# Meta-analysis of 375,000 individuals identifies 38 susceptibility loci for migraine

**DOI:** 10.1101/030288

**Authors:** Padhraig Gormley, Verneri Anttila, Bendik S Winsvold, Priit Palta, Tonu Esko, Tune H. Pers, Kai-How Farh, Ester Cuenca-Leon, Mikko Muona, Nicholas A Furlotte, Tobias Kurth, Andres Ingason, George McMahon, Lannie Ligthart, Gisela M Terwindt, Mikko Kallela, Tobias M Freilinger, Caroline Ran, Scott G Gordon, Anine H Stam, Stacy Steinberg, Guntram Borck, Markku Koiranen, Lydia Quaye, Hieab HH Adams, Terho Lehtimäki, Antti-Pekka Sarin, Juho Wedenoja, David A Hinds, Julie E Buring, Markus Schürks, Paul M Ridker, Maria Gudlaug Hrafnsdottir, Hreinn Stefansson, Susan M Ring, Jouke-Jan Hottenga, Brenda WJH Penninx, Markus Färkkilä, Ville Artto, Mari Kaunisto, Salli Vepsäläinen, Rainer Malik, Andrew C Heath, Pamela A F Madden, Nicholas G Martin, Grant W Montgomery, Eija Hämäläinen, Hailiang Huang, Andrea E Byrnes, Lude Franke, Jie Huang, Evie Stergiakouli, Phil H Lee, Cynthia Sandor, Caleb Webber, Zameel Cader, Bertram Muller-Myhsok, Stefan Schreiber, Thomas Meitinger, Johan G Eriksson, Veikko Salomaa, Kauko Heikkilä, Elizabeth Loehrer, Andre G Uitterlinden, Albert Hofman, Cornelia M van Duijn, Lynn Cherkas, Linda M. Pedersen, Audun Stubhaug, Christopher S Nielsen, Minna Männikkö, Evelin Mihailov, Lili Milani, Hartmut Göbel, Ann-Louise Esserlind, Anne Francke Christensen, Thomas Folkmann Hansen, Thomas Werge, Jaakko Kaprio, Arpo J Aromaa, Olli Raitakari, M Arfan Ikram, Tim Spector, Marjo-Riitta Järvelin, Andres Metspalu, Christian Kubisch, David P Strachan, Michel D Ferrari, Andrea C Belin, Martin Dichgans, Maija Wessman, Arn MJM van den Maagdenberg, John-Anker Zwart, Dorret I Boomsma, George Davey Smith, Kari Stefansson, Nicholas Eriksson, Mark J Daly, Benjamin M Neale, Jes Olesen, Daniel I. Chasman, Dale R Nyholt, Aarno Palotie, on behalf of the International Headache Genetics Consortium

## Abstract

Migraine is a debilitating neurological disorder affecting around 1 in 7 people worldwide, but its molecular mechanisms remain poorly understood. Some debate exists over whether migraine is a disease of vascular dysfunction, or a result of neuronal dysfunction with secondary vascular changes. Genome-wide association (GWA) studies have thus far identified 13 independent loci associated with migraine. To identify new susceptibility loci, we performed the largest genetic study of migraine to date, comprising 59,674 cases and 316,078 controls from 22 GWA studies. We identified 45 independent single nucleotide polymorphisms (SNPs) significantly associated with migraine risk (*P* < 5 × 10^−8^) that map to 38 distinct genomic loci, including 28 loci not previously reported and the first locus identified on chromosome X. Furthermore, a subset analysis for migraine without aura (MO) identified seven of the same loci as from the full sample, whereas no loci reached genome-wide significance in the migraine with aura (MA) subset. In subsequent computational analyzes, the identified loci showed enrichment for genes expressed in vascular and smooth muscle tissues, consistent with a predominant theory of migraine that highlights vascular etiologies.

---

Migraine is currently ranked as the third most common disease worldwide, with a lifetime prevalence of 15-20%, affecting up to one billion people across the globe^1,2^. It ranks as the 7^th^ most disabling of all diseases worldwide (or 1^st^ most disabling neurological disease) in terms of years of life lost to disability^1^ and is the 3^rd^ most costly neurological disorder after dementia and stroke^3^. There is an ongoing debate about whether migraine is a disease of vascular dysfunction, or a result of neuronal dysfunction with vascular changes representing downstream effects not themselves causative of migraine^4,5^. However, genetic evidence favoring one theory versus the other is lacking. At the phenotypic level, migraine is defined by diagnostic criteria from the International Headache Society^6^. There are two prevalent sub-forms: migraine without aura (MO) is characterized by recurrent attacks of moderate or severe headache associated with nausea or hypersensitivity to light and sound. Migraine with aura (MA) is characterized by transient visual and/or sensory and/or speech symptoms usually followed by a headache phase similar to MO. Some patients have both forms of attacks (MO or MA) concurrently or at different times in the disease course. Acute drug treatments are given to abort a migraine attack and daily prophylactic treatments are available to reduce the number of attacks but lack of efficacy or side effects often limit their use^7^. Identifying new molecular targets or mechanisms could lead to new treatments that are more specific and more effective with fewer side effects.

Family and twin studies estimate a heritability of 42% (95% confidence interval [CI] = 36-47%) for migraine^8^, pointing to a strong genetic component of the disease. Higher estimates have been observed for the migraine subtypes, with up to 61% (95%CI = 49-71%) for MO^9,10^ and 65% (95%CI = 49-78%) for MA^10-12^. Despite this, genetic association studies have revealed relatively little about the molecular mechanisms that contribute to pathophysiology. Understanding has been limited partly because, to date, only 13 genome-wide significant risk loci have been identified for the prevalent forms of migraine^13-16^. In familial hemiplegic migraine (FHM), a rare Mendelian form of the disease, three ion transport-related genes (*CACNA1A, ATP1A2* and *SCN1A*) have been implicated^17-19^. These findings suggest that mechanisms that regulate neuronal ion homeostasis might also be involved in MO and MA, however, no genes related to ion transport have yet been identified for these more prevalent forms of migraine^20^.

We performed a meta-analysis of 22 genome-wide association (GWA) studies, consisting of 59,674 cases and 316,078 controls collected from six tertiary headache clinics and 27 population-based cohorts through our worldwide collaboration in the International Headache Genetics Consortium (IHGC). This combined dataset contained over 35,000 new migraine cases not included in previously published GWA studies. Here we present the findings of this new meta-analysis, including 38 genomic loci, harboring 45 independent association signals identified at levels of genome-wide significance, which support current theories of migraine pathophysiology and also offer new insights into the disease.

## Results

### Significant associations at 38 independent genomic loci

The primary meta-analysis was performed on all migraine samples available through the IHGC, regardless of ascertainment. These case samples include both individuals diagnosed with migraine by a doctor as well as individuals with self-reported migraine via questionnaires. Study design and sample ascertainment for each individual study is outlined in the supplementary material (**Supplementary Methods and Supplementary Table 1**). The final combined sample for the main analysis consisted of 59,674 cases and 316,078 controls in 22 non-overlapping case-control samples (**Table 1**). All samples were of European ancestry.

**Table 1.**
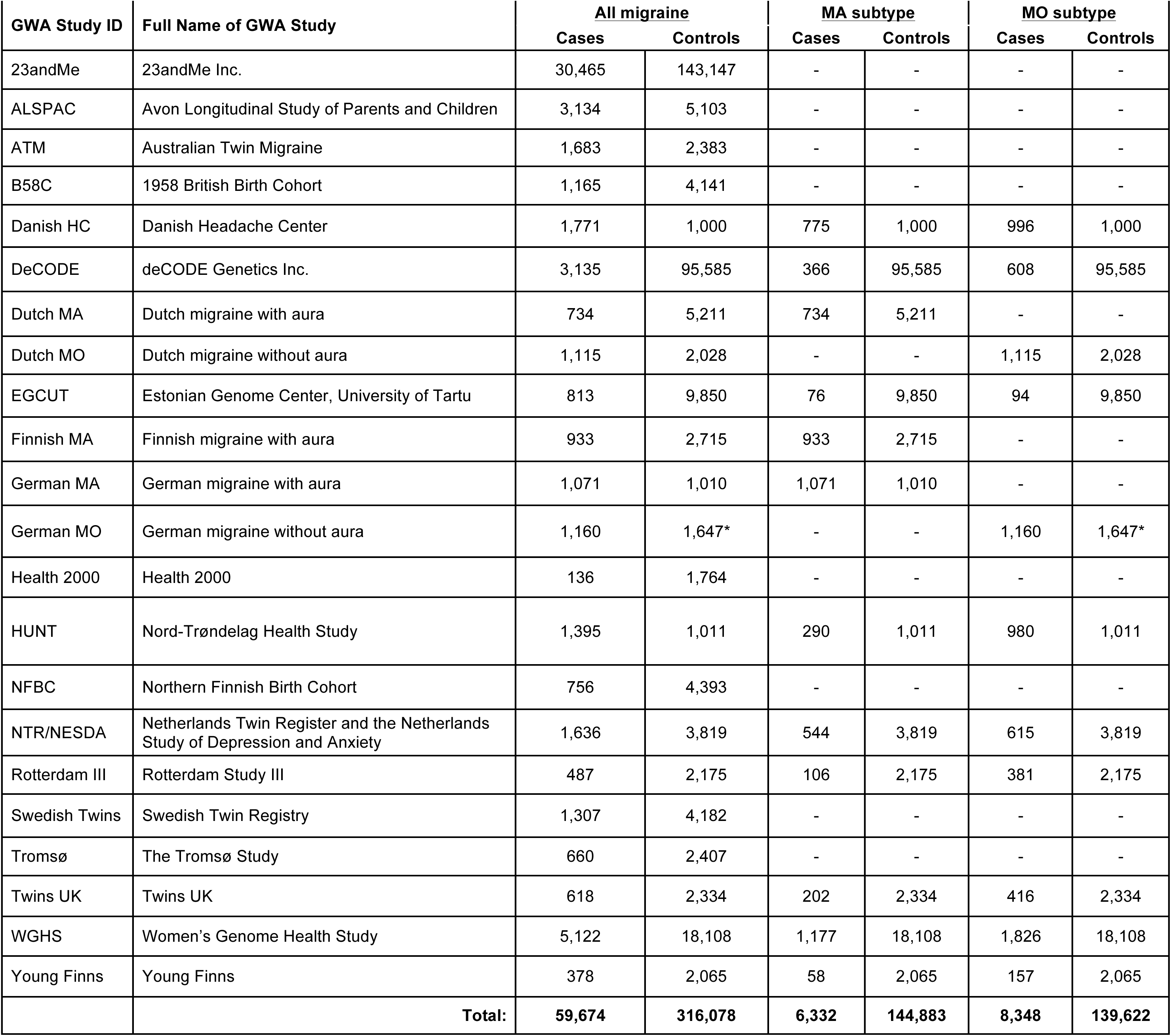
Individual IHGC GWA studies listed with cases and control numbers used in the primary analysis (all migraine) and in the sub-analyzes (MA and MO subtypes). Note that chromosome X genotype data was unavailable from three of the individual GWA studies (EGCUT, Rotterdam III, and TwinsUK) and also one of the German MO control studies (GSK) meaning that the number of samples analyzed on chromosome X was 57,756 cases and 299,109 controls. Complete data was available on the autosomes for all samples.

The 22 individual GWA studies completed standard quality control protocols summarized in **Supplementary Table 2.** After quality control, missing genotypes (SNPs and short insertion-deletions) were imputed into each sample using a common 1000 Genomes Project reference panel (Phase I, v3, March 2012 release)^21^. Association analyzes were performed within each sample using logistic regression on the imputed marker dosages while adjusting for sex and principal components (when necessary) to adjust for sub-European population structure. These association results were combined using an inverse-variance weighted fixed-effects meta-analysis. Markers included in the final meta-analysis were also filtered based on individual study quality metrics (imputation INFO score ≥ 0.6, MAF ≥ 0.01) and combined quality metrics (heterogeneity index *i*^2^ < 0.75 and successfully genotyped/imputed in at least half of the 22 GWA studies). This left 8,094,889 variants for consideration in our primary analysis.

**Table 2.**
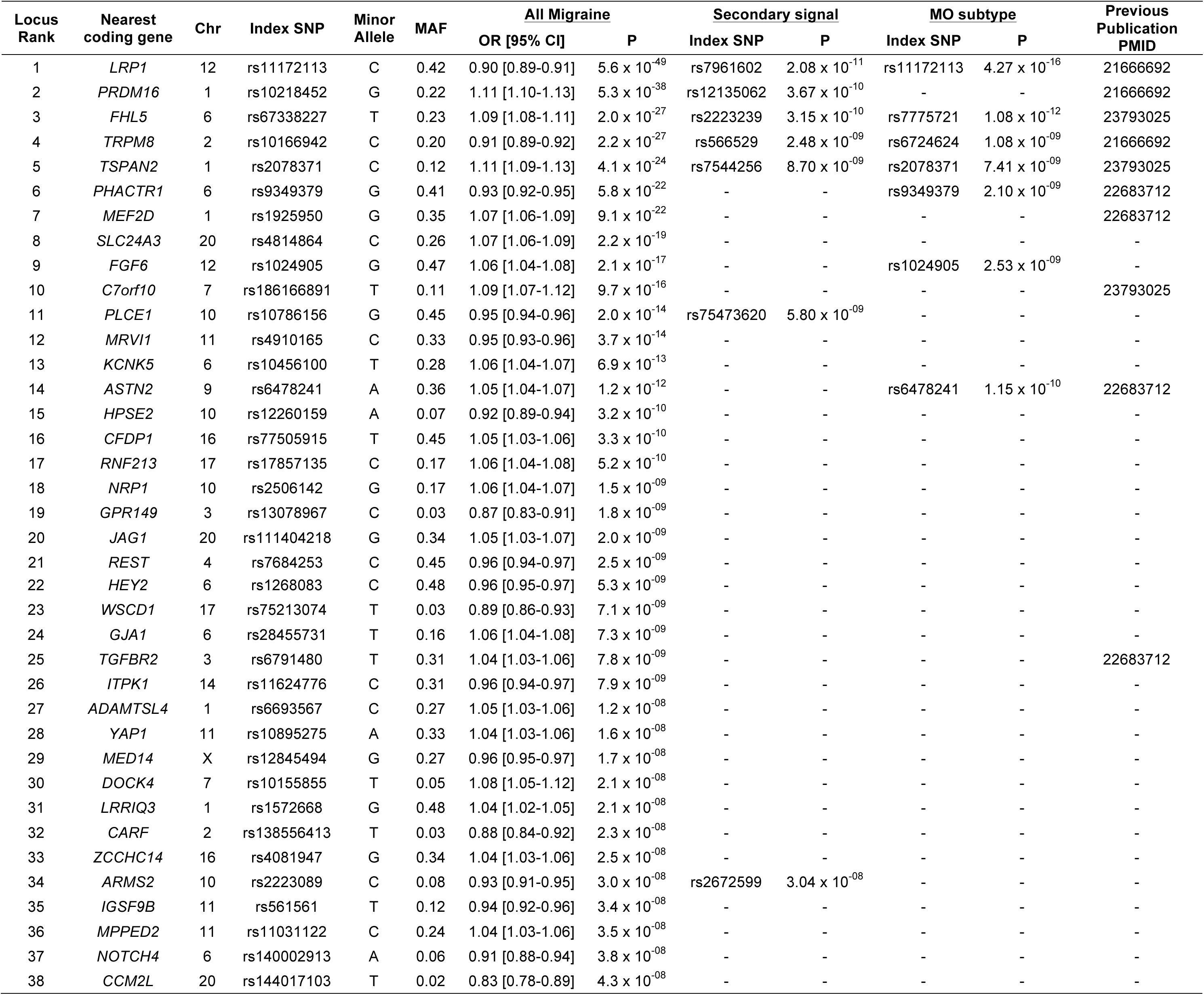
Summary of the 38 genomic loci associated with the prevalent types of migraine. Ten loci were previously reported (PubMed IDs listed) and 28 are newly found in this study. For each locus, the nearest coding gene to the index SNP is given. For loci that contain a secondary LD-independent signal passing genome-wide significance, the secondary index SNP and P-value is given. For the seven loci reaching genome-wide significance in the MO sub-type analysis, the corresponding index SNP and P-value are also given.

Among these variants in the primary analysis sample, we identified 45 genome-wide significant SNP associations (*P* < 5 × 10^−8^) that are independent (*r*^2^ < 0.1) with regards to linkage disequilibrium (LD). To help identify candidate risk genes from these, we defined an associated locus as the genomic region bounded by all markers in LD (*r*^2^ > 0.6 in 1000 Genomes, Phase I, EUR individuals) with each of the 45 index SNPs and in addition, all such regions in close proximity (< 250 kb) were merged. From these defined regions we implicate 38 distinct genomic loci in total for the prevalent forms of migraine, 28 of which have not previously been reported (see **Figure 1**), including the first genome-wide associated locus for migraine on chromosome X.

**Figure 1.**
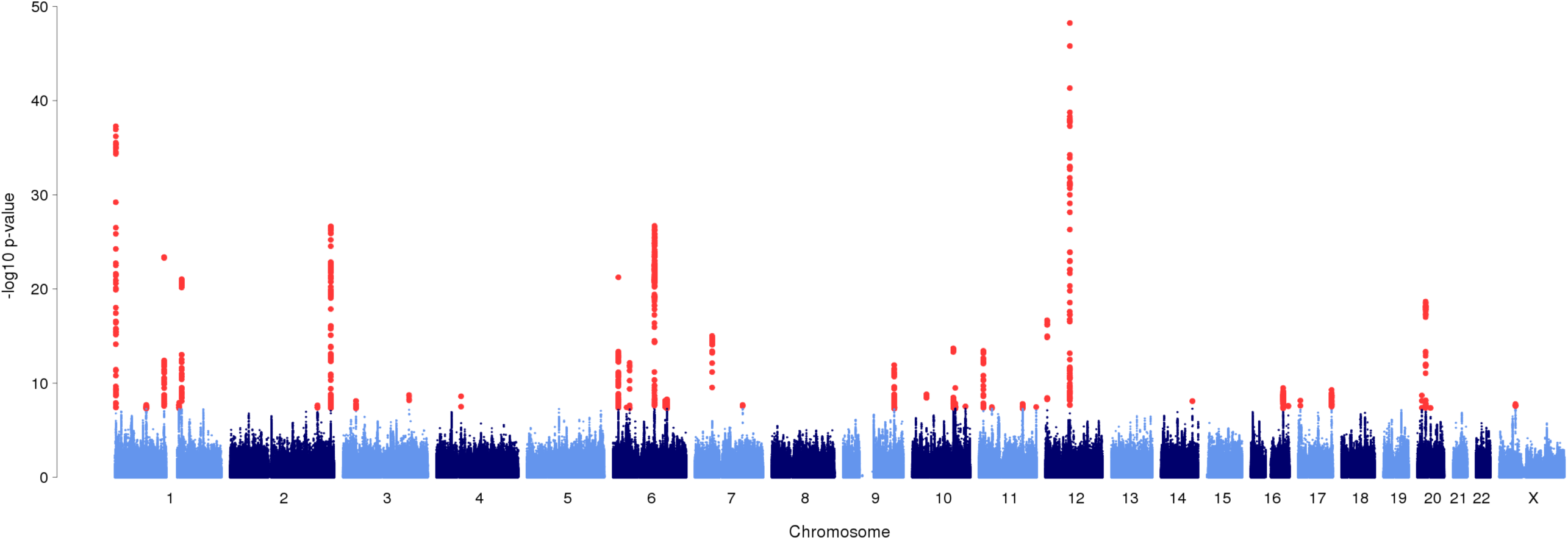
Manhattan plot from the primary meta-analysis of all migraine (59,674 cases *vs.* 316,078 controls). A fixed-effects meta-analysis was used to combine the association statistics from all 22 clinic and population-based studies from the IHGC. The x-axis shows the chromosomal position and the y-axis shows the significance of tested markers from logistic regression. Markers with test statistics that reach genome-wide significance (*P* < 5 × 10^−8^) are shown in red.

These 38 loci replicate 10 of the 13 previously reported genome-wide associations to migraine (see **Table 2** for a summary of the loci identified). At seven of the 10 replicated known loci we now find a more significant index SNP (rs4379368 to rs186166891 at *C7orf10,* rs13208321 to rs67338227 at *FHL5,* rs6790925 to rs6791480 at *TGFBR2,* rs7577262 to rs10166942 at *TRPM8,* rs2274316 to rs1925950 at *MEF2D,* rs12134493 to rs2078371 at *TSPAN2,* and rs2651899 to rs10218452 at *PRDM16*). Seven of the 38 loci contain a secondary genome-wide significant SNP (*P* < 5 × 10^−8^) that is not in LD (*r*^2^ < 0.1) with the top SNP in the locus (**Table 2**). Five of these secondary signals were found in known loci (at *LRP1, PRDM16, FHL5, TRPM8,* and *TSPAN2*), while the others were found within two of the 28 new loci (*PLCE1* and *ARMS2*). Therefore, out of the 45 LD-independent SNPs reported here, 35 are new associations to migraine. Three previously reported loci that were associated to subtypes of migraine (rs1835740 near *MTDH,* rs10915437 near *AJAP1,* and rs10504861 near *MMP16*)^13,16^ show only nominal significance (*P* < 5 × 10^−3^) in the current meta-analysis (**Supplementary Table 3**), however, these loci have since been shown to be associated to specific phenotypic features of migraine^22^ and therefore may require a more phenotypically homogeneous sample to be accurately assessed for association with migraine. Most of the effects at the 45 identified index SNPs were homogeneous, but four SNPs (at *TRPM8, ZCCHC14, MRVI1,* and *CCM2L*) exhibited some moderate heterogeneity across the individual GWA studies (Cochran’s Q test *p-value* < 0.05, **Supplementary Table 4**).

### Characterization of the associated loci

In total, 32 of 38 (84%) genomic loci overlap with transcripts from protein-coding genes, and 17 (45%) of these regions contain just a single gene (see **Supplementary Figure 1** for regional plots of the 38 genomic loci and **Supplementary Table 5** for extended information on each locus). Among the 38 genomic loci, only two contain ion channel genes (*KCNK5*^23^ and *TRPM8*^24^). Hence, despite previous hypotheses of migraine as a potential channelopathy^5,25^, the loci identified to date do not support common variants in ion channel genes as strong susceptibility components in prevalent forms of migraine. However, three other loci do contain genes involved more generally in ion homeostasis (*SLC24A3*^26^, *ITPK1*^27^, and GJA1^28^, **Supplementary Table 6**).

Several of the genes have previous associations to vascular disease (*PHACTR1*,^29,30^ *TGFBR2*,^31^ *LRP1*,^32^ *PRDM16*,^33^ *RNF213*,^34^ *JAG1*,^35^ *HEY2*,^36^ *GJA1*^37^, *ARMS2*^38^), or are involved in smooth muscle contractility and regulation of vascular tone (*MRVI1,*^39^ *GJA1*,^40^ *SLC24A3*,^41^ *NRP1*^42^). Two of the 45 migraine SNPs have previously reported associations in the National Human Genome Research Institute (NHGRI) GWAS catalog at exactly the same SNP (rs9349379 at *PHACTR1* with coronary heart disease^43–45^ and coronary artery calcification^46^ and rs11624776 at *ITPK1* with thyroid hormone levels^47^). Six of the loci harbor genes that are involved in nitric oxide signaling and oxidative stress (*REST*^48^, *GJA1*^49^, *YAP1*^50^, *PRDM16*^51^, *LRP1*^52^, and *MRVI1*^53^).

From each locus we chose the nearest gene to the index SNP to assess gene expression activity in various tissues from the GTEx consortium (**Supplementary Figure 2**). While we found that most of the putative genes in the migraine loci were expressed in many different tissue types (including brain and vascular tissues), we could detect tissue specificity in certain instances whereby some genes appear to be more active in one particular tissue group relative to the others. For instance six genes were relatively more actively expressed in brain (*ASTN2, CFDP1, DOCK4, ITPK1, MPPED2,* and *WSCD1*) compared to other tissues, whereas two genes were specifically active in vascular tissues (*HEY2* and *PRDM16*). Many of the other putative genes in the migraine loci were actively expressed in more than one tissue group.

**Figure 2.**
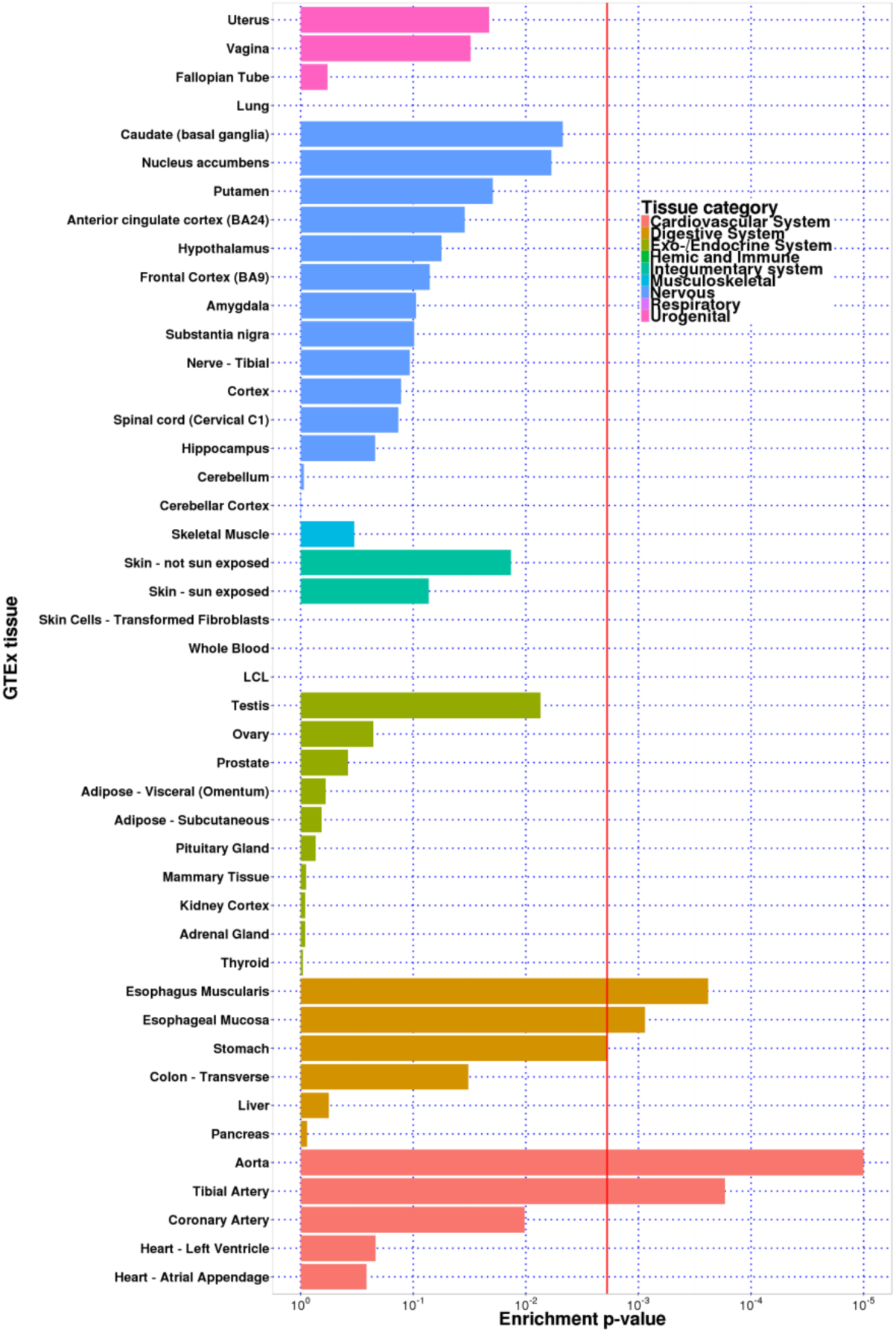
Gene expression enrichment in specific tissues using genes at the 38 migraine loci and RNAseq data from the GTEx consortium.

### Genomic inflation and LD-score regression analysis

To assess whether the 38 independent genomic loci harbor true associations with migraine rather than reflecting systematic differences between cases and controls (such as population stratification) we analyzed the genome-wide inflation of test statistics in our primary meta-analysis dataset. As expected for a complex polygenic trait, the distribution of test statistics deviates from the null (genomic inflation factor *λ_GC_* = 1.24, **Supplementary Figure 3**) which is in line with other large GWA study meta-analyzes^54-57^. Since much of the inflation in a polygenic trait arises from LD between the causal SNPs and many other neighboring SNPs in the local region, we LD-pruned the meta-analysis results to create a set of essentially LD-independent markers. The LD-pruning was performed in PLINK^58^ using a sliding window (size 250-kb) that removes one marker from every pair that is in LD (*r*^2^ > 0.2). The resulting genomic inflation was reduced (*λ_GC_* = 1.15, **Supplementary Figure 4**) and reflects the polygenic signal remaining from independent loci.

**Figure 3.**
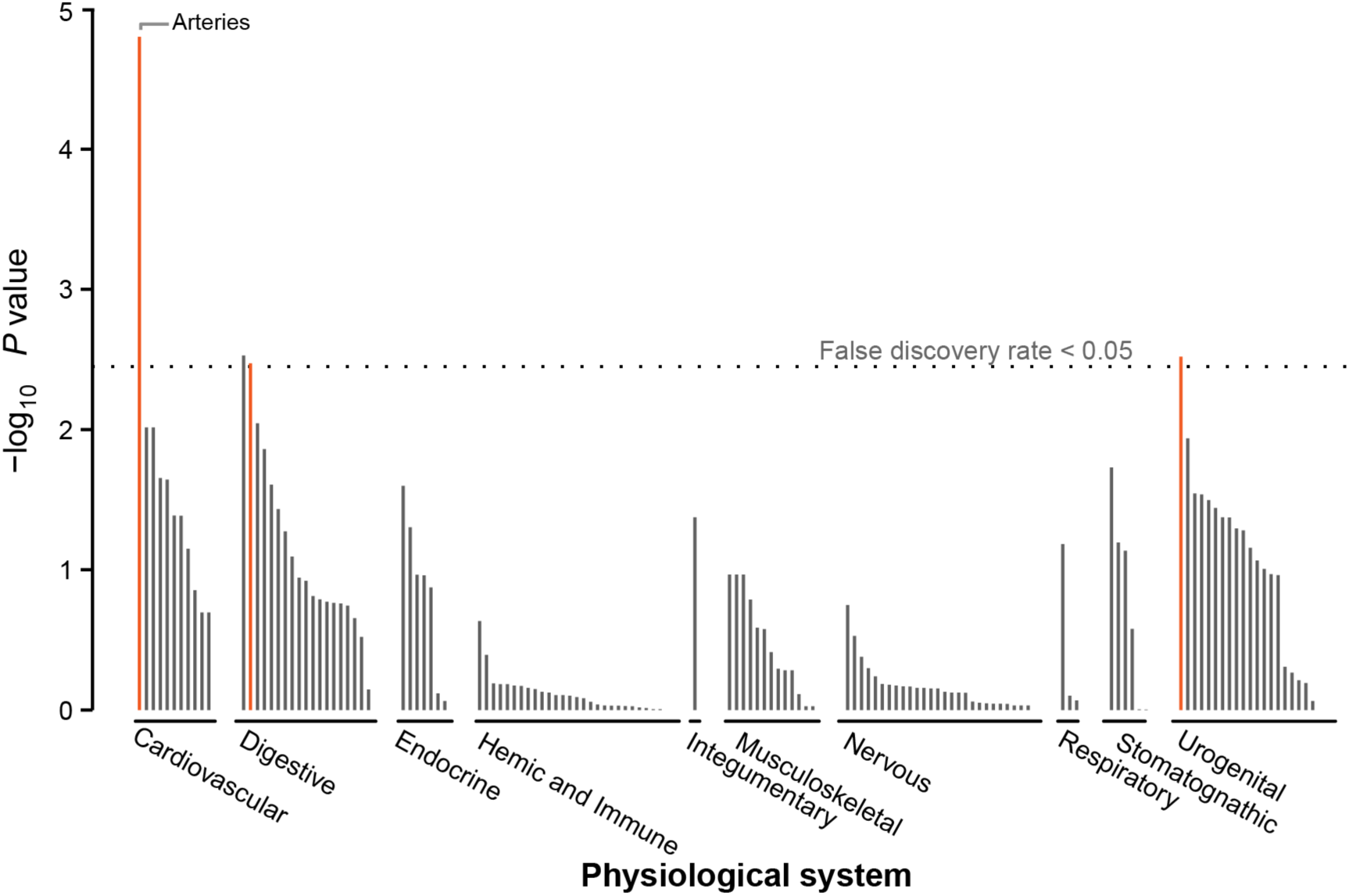
Gene expression enrichment in various tissues of the 38 migraine loci using microarray data from DEPICT.

**Figure 4.**
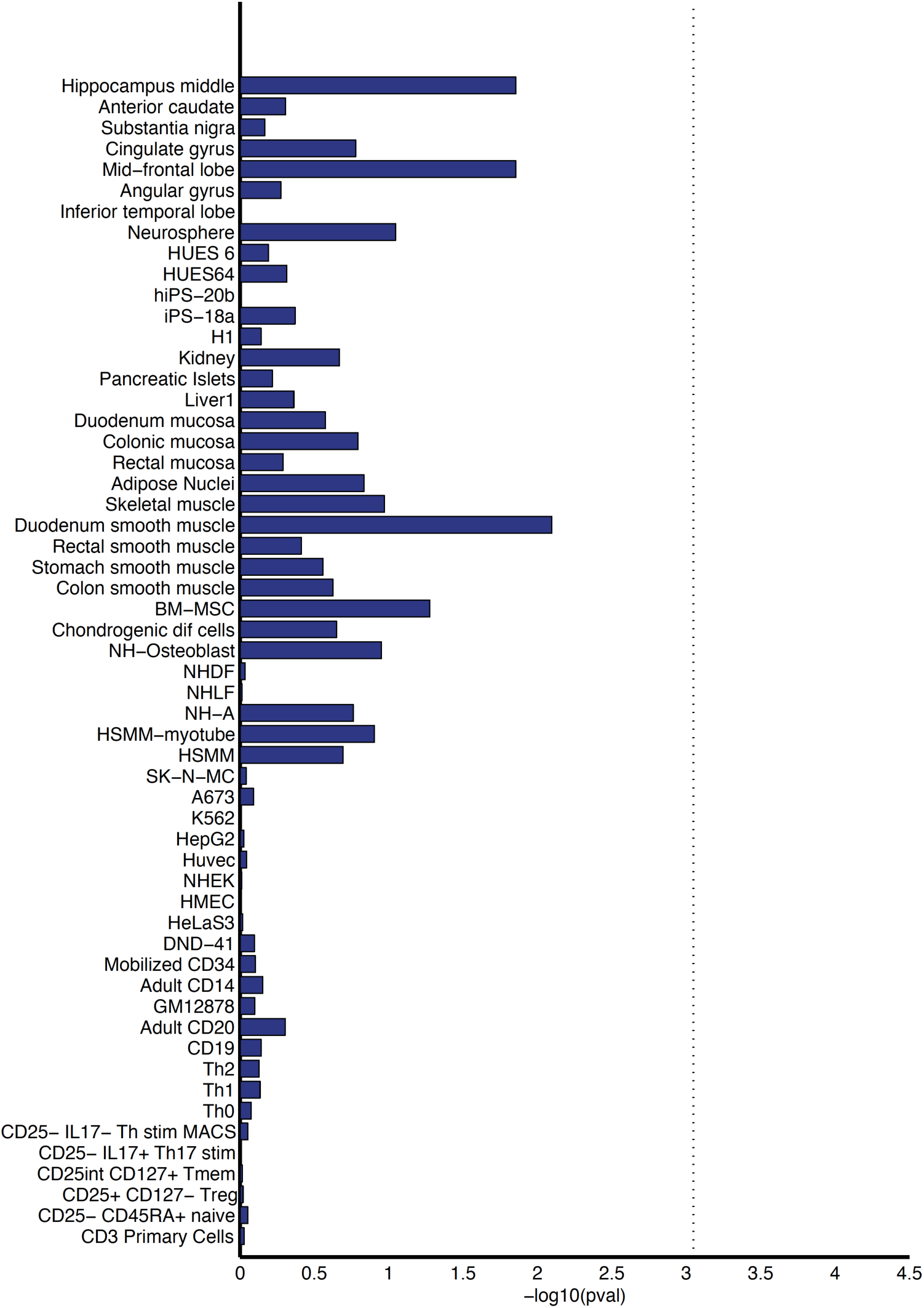
Enrichment of the migraine loci in sets of tissue-specific enhancers. We mapped credible sets from the 38 genome-wide significant loci to sets of enhancers under active expression in 56 different tissues and cell lines (identified by enrichment of H3K27ac histone marks in the roadmapepigenomics.org data). The dashed line represents the Bonferroni-corrected p-value threshold for 56 separate test

In order to confirm that the deviation between our observed test statistics and the null distribution is primarily coming from true polygenic signal, we analyzed our meta-analysis results using LD-score regression^59^. This method tests for a linear relationship between marker test statistics and LD score, defined as the sum of *r*^2^ values between a marker and all other markers within a 1-Mb window. The primary analysis results show a linear relationship between association test statistics and LD-score (**Supplementary Figure 5**) suggesting that the deviation in test statistics consists mainly of true polygenic signal rather than population stratification or other confounders. These results are consistent with the theory of polygenic disease architecture shown previously by both simulation and real data for GWAS samples of similar size^60^.

**Figure 5.**
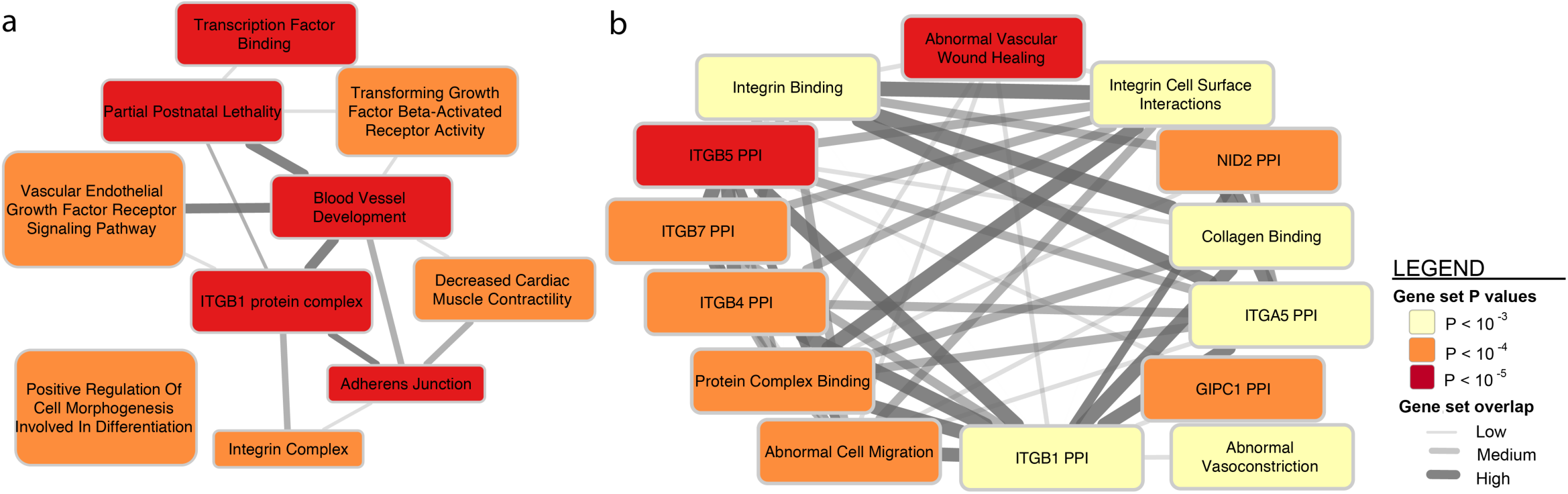
DEPICT network of the reconstituted gene sets that were significantly enriched (*FDR* < 0.05) for genes at the migraine loci. **a)** Shows 10 groups of the reconstituted gene sets clustered by similarity. **b)** Shows an example of the significant gene sets inside the *ITGB1 PPI* cluster. A full list of the 67 significantly enriched gene sets can be found in **Supplementary Table 12.**

### MA and MO subtype analyzes

To elucidate the pathophysiological mechanisms underpinning the migraine aura, we performed a secondary analysis of the data by creating two subsets that included only samples with the subtypes MA and MO. These subsets only included those studies where sufficient information was available to assign a diagnosis of either subtype according to classification criteria standardized by the International Headache Society (IHS). Specific IHS diagnostic criteria used to classify MA and MO can be found in the 2^nd^ edition of the International Classification of Headache Disorders (ICHD-II)^6^. For the population-based study samples this involved questionnaires to assess the necessary criteria, whereas for the clinic-based study samples the diagnosis was assigned on the basis of a structured interview by telephone or in person. A stricter diagnosis is required for these migraine subtypes as the migraine aura specifically is challenging to distinguish from other neurological features that can present as symptoms from unrelated conditions.

As a result, the migraine subtype analyzes consisted of considerably smaller sample sizes compared to the main analysis (6,332 cases *vs.* 144,883 controls for MA and 8,348 cases *vs.* 139,622 controls for MO, see **Table 1**). As with the primary migraine analysis, the signals in the test statistics for MA or MO were consistent with underlying polygenic architecture rather than other potential sources of inflation (**Supplementary Figure 6 and 7**). For the MO subset analysis we found seven independent genomic loci (near *TSPAN2, TRPM8, PHACTR1, FHL5, ASTN2,* near *FGF6,* and *LRP1*) to be significantly associated with MO (**Table 2** and **Supplementary Figure 8**). All seven of these loci were already identified in our primary analysis of ‘all migraine’ types, possibly reflecting the fact that MO is the most common form of migraine (around 2 in 3 cases) and likely drives the association signals in the primary analysis. Notably, no loci were associated to migraine with aura in the MA subset analysis (**Supplementary Figure 9**).

To investigate whether excess heterogeneity could be contributing to the lack of associations in MA, we performed a heterogeneity analysis between the two subgroups. First we created two subsets of the MA and MO datasets from which none of the case or control individuals were overlapping (**Supplementary Table 7**). Then we selected the 45 LD-independent SNPs associated from the primary analysis and used a random-effects model to combine the MA and MO samples in a meta-analysis that allows for heterogeneity between the two migraine groups^61^. We found little heterogeneity with only seven of the 45 SNPs (at *REST, MPPED2, PHACTR1, ASTN2, MEF2D, PLCE1*, and *MED14*) exhibiting some signs of heterogeneity between the two groups (**Supplementary Table 8**).

### Credible sets of markers within each locus

For each of the 38 migraine-associated loci, we defined a credible set of markers that could plausibly be considered as causal using a Bayesian-likelihood based approach^62^. This method incorporates evidence from association test statistics and the LD structure between SNPs in a locus (for a full description, see **Supplementary Methods**). We found three instances (in the *RNF213, PLCE1,* and *MRVI1* loci) where the association signal could be credibly attributed to exonic missense polymorphisms (**Supplementary Table 9**). However, most of the credible markers at each locus were either intronic or intergenic, which is consistent with the theory that most variants detected by GWA studies involve regulatory effects on gene expression rather than disrupting protein structure^63,64^.

### Overlap with eQTLs in brain and blood

To try to identify specific migraine loci that might influence gene expression, we used previously published datasets that catalog expression quantitative trait loci (eQTLs) in either of two studies from peripheral venous blood (N_1_ = 3,754 and N_2_ = 2,360) or a third study from human brain cortex tissue (N_3_ = 550). Using these datasets we applied two methods to identify eQTLs that could explain associations at the 38 migraine loci (**Supplementary Methods**). The first approach tested whether the migraine index SNP at each locus was a significant cis-eQTL after conditioning on the best local eQTL to transcripts within a 1-Mb window (**Supplementary Table 10**). The second more stringent approach merged the migraine credible sets defined above with credible sets from cis-eQTL signals within a 1-Mb window and tested if the association signals between the migraine and eQTL credible sets were correlated (**Supplementary Tables 11-12 and Supplementary Figure 10**). After adjusting for multiple testing we found only two plausible eQTL associations in peripheral blood, rs7684253 (at the *REST* locus) as an eQTL to *NOA1* and rs6693567 (at the *ADAMTSL4* locus) as an eQTL to *MLLT11* (**Supplementary Table 10**). This low number (2 out of 38) is consistent with previous studies which have observed that available eQTL catalogues currently lack sufficient tissue specificity and developmental diversity to provide enough power to provide meaningful biological insight^56^. No plausibly causal eQTLs were observed in expression data from brain.

### Gene expression enrichment in specific tissues

To understand if the 38 migraine loci as a group are enriched for expression in certain tissue groups, we analyzed RNA-seq data (from 1,641 samples in 175 individuals across 42 tissues and three cell lines) generated as part of the pilot phase of the Genotype-Tissue Expression (GTEx) project^65^. Using this data on gene expression, we tested whether genes near to credibly causal SNPs at the 38 migraine loci were significantly enriched in certain tissues (**Supplementary Methods**). We found four tissues that were significantly enriched (after Bonferroni correction) for expression of the migraine genes (**Figure 2**). The two most strongly enriched tissues were part of the cardiovascular system; the *aorta* and *tibial artery.* Two other significant tissues were from the digestive system; *esophagus muscularis* and *esophageal mucosa.* We replicated these enrichment results in an independent dataset using a component of the DEPICT^66^ tool that conducts a tissue-specific enrichment analysis on microarray-based gene expression data (**Supplementary Methods**). DEPICT highlighted four tissues (**Figure 3 and Supplementary Table 13**) with significant enrichment of the genes within the migraine loci; arteries (*P* = 1.58 × 10^−5^), the upper gastrointestinal tract (*P* = 2.97 × 10^−3^), myometrium (*P* = 3.03 × 10^−3^), and stomach (*P* = 3.38 × 10^−3^).

Taken together, the expression analyzes implicate arterial and gastrointestinal (GI) tissues. To try to determine whether the enrichment signature of arterial and gastrointestinal tissues could be attributed to a more specific type of smooth muscle, we examined the expression of the nearest protein-coding genes at migraine loci in a panel of 60 different types of human smooth muscle tissue drawn from gastrointestinal, genitourinary, arterial, venous, and bronchial sources^67^. Overall, migraine loci genes were not significantly enriched in a particular class of smooth muscle, although anecdotal examples such as *TGFBR2* demonstrated striking vascular expression (**Supplementary Figures 11-13**). This suggests that the enrichment of migraine disease variants in genes expressed in tissues with a smooth muscle component is not specific to blood vessels, the stomach or GI tract, but rather appears to be generalizable across vascular and visceral smooth muscle types. Future studies will be required to be able to identify any specific tissues that are involved.

Combined, these results suggest that some of the genes affected by migraine-associated variants are highly expressed in vascular tissues and their dysfunction could play a role in migraine. Furthermore, the enrichment results suggest that other tissue types (e.g. smooth muscle) could also play a role and this may become evident once more migraine loci are discovered.

### Enrichment in tissue-specific enhancers

To assess, from a different perspective, the hypothesis that migraine variants might operate via effects on gene-regulation, we investigated the degree of overlap with histone modifications. We identified candidate causal variants underlying the 38 migraine loci, and examined their enrichment within cell-type specific enhancers from 56 primary human tissues and cell types from the Roadmap Epigenomics^68^ and ENCODE projects^69^ (**Supplementary Methods**). Candidate causal variants showed highest enrichment in tissues from the mid-frontal lobe and duodenum smooth muscle, but these enrichments were not significant after adjusting for multiple testing (**Figure 4**).

### Gene set enrichment analyzes

To implicate underlying biological pathways involved in migraine, we applied a Gene Ontology (GO) over-representation analysis of the 38 migraine loci (**Supplementary Methods**). We found nine vascular-related biological function categories that are significantly enriched after correction for multiple testing, including *circulatory system development* (GO:0072359) and *blood vessel development* (GO:0001568) (**Supplementary Table 14**). Interestingly, we found little statistical support from the identified loci for some molecular processes that have been previously linked to migraine, e.g. ion homeostasis, glutamate signaling, serotonin signaling, nitric oxide signaling, and oxidative stress (**Supplementary Table 15**). However, it is possible that the lack of enrichment for these functions may be explained by recognizing that current annotations for many genes and pathways are still far from comprehensive, or that larger numbers of migraine loci need to be identified before we have sensitivity to detect enrichment in these mechanisms.

For a comprehensive pathway analysis tool we used DEPICT, which incorporates co-expression information from gene expression microarray data to implicate additional, functionally less well-characterized genes in known biological pathways, protein-protein complexes and mouse phenotypes^66^ (by forming so-called ‘reconstituted gene sets’). From DEPICT we identified 67 reconstituted gene sets that are significantly enriched (FDR < 5%) for genes found among the 38 migraine associated loci (**Supplementary Table 16**). Because the reconstituted gene sets had genes in common, we clustered them into 10 distinct groups of gene sets (**Figure 5 and Supplementary Methods**). Several gene sets, including the most significantly enriched reconstituted gene set *(Abnormal Vascular Wound Healing; P* = 1.86 × 10^−6^), were grouped into clusters related to cell-cell interactions *(ITGB1 PPI, Adherens Junction, Integrin Complex).* Several of the other gene set clusters were related to vascular-biology *(Blood Vessel Development, Cellular Response To Vascular Endothelial Growth Factor Stimulus,* **Figure 5 and Supplementary Table 16**).

## Discussion

In what is the largest genetic study of migraine to date, we identified 38 distinct genomic loci harboring 45 independent susceptibility markers for the prevalent forms of migraine. We provide evidence that migraine-associated genes identified through variants of small effect are involved both in arterial and smooth muscle function. Two separate analyzes, the DEPICT and the GTEx gene-expression enrichment analyzes, together point to vascular and smooth muscle tissues being involved in common variant susceptibility to migraine. The vascular finding is consistent with known co-morbidities and previously reported shared polygenic risk between migraine, stroke and cardiovascular diseases^70,71^. Furthermore, a recent GWA study of Cervical Artery Dissection (CeAD) identified a genome-wide significant association at exactly the same index SNP (rs9349379) as is associated to migraine in the *PHACTR1* locus, suggesting the possibility of partially shared genetic components between migraine and CeAD^30^. These results suggest that vascular dysfunction and possibly also other smooth muscle dysfunction likely play roles in migraine pathogenesis.

The support for vascular and smooth muscle enrichment of the loci is strong, with multiple lines of evidence from independent methods and independent datasets. However, it remains likely that neurogenic mechanisms are also involved in migraine. For example, several lines of evidence from previous studies have pointed to neurogenic mechanisms in migraine^5,72-75^. We found some support for this when looking at gene expression of individual genes at the 38 loci (**Supplementary Figure 2**), where many specific genes were active in brain tissues and therefore could have neuronal function. While we did not observe statistically significant enrichment in brain across all loci, it may be that more associated loci are needed to detect this. Alternatively, it could be due to difficulties in collecting appropriate brain tissue samples with enough specificity, or other technical challenges. An additional limiting factor is that there is less clarity of the biological mechanisms for a brain disease like migraine compared to some other common diseases, e.g. autoimmune diseases or cardio-metabolic diseases where intermediate risk factors and underlying mechanisms are better understood.

Interestingly, some of the analyzes highlight gastrointestinal tissues. Although migraine attacks may include gastrointestinal symptoms (e.g. nausea, vomiting, diarrhea)^76^ it is likely that the signals observed here broadly represent smooth muscle signals rather than gastrointestinal specificity. Smooth muscle is a predominant tissue of the intestine, yet specific smooth muscle subtypes were not available to test this hypothesis in our primary enrichment analyzes. We showed instead in a range of 60 smooth muscle subtypes, that the migraine loci are expressed in many types of smooth muscle, including vascular (**Supplementary Figure 12 and 13**). These results, while not conclusive, suggest that the enrichment of the migraine loci in smooth muscle is not specific to the stomach and GI tract.

Our results identify specific cellular pathways and provide an opportunity to determine whether the genomic data supports previously presented hypotheses of pathways linked to migraine. One prevailing hypothesis stimulated by findings in familial hemiplegic migraine (FHM) has been that migraine is a channelopathy^5,25^. Among the 38 migraine loci only two harbor known ion channels (*KCNK5*^23^ and *TRPM8*^24^), while three additional loci (*SLC24A3*^26^, *ITPK1*^27^, and *GJA1*^28^) can be linked to ion homeostasis. This further supports the findings of previous studies that in common forms of migraine, ion channel dysfunction is not the major pathophysiological mechanism^20^. However, more generally, genes involved in ion homeostasis could be a component of the genetic susceptibility. Moreover, we cannot exclude that ion channels could still be important contributors in MA, the form most closely resembling FHM, as our ability to identify loci in this subgroup is more challenging. Another suggested hypothesis relates to oxidative stress and nitric oxide (NO) signaling. Nitroglycerine and sodium nitroprusside, two pro-drugs delivering NO to tissues can both induce headache in normal patients and migraine in migraine patients^77,78^. Animal experiments have shown relations to superoxide and oxidative stress^79^. Six genes with known links to oxidative stress and NO, within these 38 loci were identified (*REST*^48^, *GJA1*^49^, *YAP1*^50^, *PRDM16*^51^, *LRP1*^52^, and *MRVI1*^53^). This is in line with previous findings^16^, however, in the DEPICT pathway analysis we found that NO-related reconstituted gene sets were not significantly associated with migraine (*FDR* > 0.54) (**Supplementary Table 15**).

In conclusion, the 38 genomic loci identified in this study support the notion that factors in vascular and smooth muscle tissues contribute to migraine pathophysiology and that the two major subtypes of migraine, MO and MA, have a partially shared underlying genetic susceptibility profile.

## URLs

1000 Genomes Project, http://www.1000genomes.org/; BEAGLE,

http://faculty.washington.edu/browning/beagle/beagle.html; DEPICT,

www.broadinstitute.org/mpg/depict; GTEx,

www.gtexportal.org; GWAMA,

http://www.well.ox.ac.uk/gwama/; IMPUTE2,

https://mathgen.stats.ox.ac.uk/impute/imputev2.html; International Headache Genetics Consortium, http://www.headachegenetics.org/; MACH,

http://www.sph.umich.edu/csg/abecasis/MACH/tour/imputation.html; matSpD,

http://neurogenetics.qimrberghofer.edu.au/matSpD; PLINK,

http://pngu.mgh.harvard.edu/~purcell/plink/; ProbABEL,

http://www.genabel.org/packages/ProbABEL; Roadmap Epigenomics Project,

http://www.roadmapepigenomics.org/; SHAPEIT,

http://mathgen.stats.ox.ac.uk/geneticssoftware/shapeit/shapeit.v778.html; SNPTEST,

https://mathgen.stats.ox.ac.uk/genetics_software/snptest/snptest.html.

## Acknowledgments

We would like to thank the numerous individuals who contributed to sample collection, storage, handling, phenotyping and genotyping within each of the individual cohorts. We also thank the important contribution to research made by the study participants. We are grateful to Huiying Zhao (QIMR Berghofer Medical Research Institute) for helpful correspondence on the pathway analyzes. We acknowledge the support and contribution of pilot data from the GTEx consortium. A list of study-specific acknowledgements can be found in the Supplementary Information.

## Author Contributions

All authors contributed to the final version of the manuscript.

